# A Nodal/Eph signalling relay drives the transition from apical constriction to apico-basal shortening in ascidian endoderm invagination

**DOI:** 10.1101/418988

**Authors:** Ulla-Maj Fiuza, Takefumi Negishi, Alice Rouan, Hitoyoshi Yasuo, Patrick Lemaire

**Affiliations:** CRBM, University of Montpellier, CNRS, Montpellier, France; Laboratoire de Biologie du Développement de Villefranche-sur-mer, CNRS, Sorbonne Universités, 06230 Villefranche-sur-mer, France

**Keywords:** Nodal, Ephrin, Gastrulation, morphogenesis, cell fate

## Abstract

Gastrulation is the first major morphogenetic event during animal embryogenesis. Ascidian gastrulation starts with the invagination of 10 endodermal precursor cells between the 64- and late 112-cell stages. This process occurs in the absence of endodermal cell division and in two steps, driven by myosin-dependent contractions of the acto-myosin network. First, endoderm precursors constrict their apex. Second, they shorten apico-basally, while retaining small apical surfaces, thereby causing invagination. The mechanisms controlling the endoderm mitotic delay, the step 1 to step 2 transition, and apico-basal shortening have remained elusive. Here, we demonstrate the conserved role during invagination of Nodal and Eph signalling in two distantly related ascidian species (*Phallusia mammillata* and *Ciona intestinalis*). We show that the transition to step 2 is controlled by Nodal relayed by Eph signalling and that Eph signalling has a Nodal-independent role in mitotic delay. Interestingly, both Nodal and Eph signals are dispensable for endodermal germ layer fate specification.

**Summary statement:** Identification of a regulatory developmental signalling sub-network driving endoderm cell shape changes during ascidian endoderm invagination, not involved in cell fate specification.

## Introduction

Epithelial invagination, the buckling of a sheet of cells into a cup-like structure, is a morphogenetic mechanism driving dramatic tissue shape changes in multiple embryonic contexts including neural tube and optic cup formation, or gastrulation. In most metazoans, endoderm invagination is the first event of gastrulation, a key embryonic process during which the embryo body plan and the main tissue types become specified (Leptin, 2005; Solnica-Krezel and Sepich, 2012). While the precise coordination of cell fate decisions, cell shape changes, cell divisions and cell movements is crucial to ensure successful embryogenesis, we only have a fragmented understanding of the way these processes are integrated at the transcriptional level (Reviewed in Heisenberg and Solnica-Krezel, 2008). In this study, we explore the mechanisms controlling endoderm invagination in the solitary ascidians *Phallusia mammillata* and *Ciona intestinalis*.

Although they diverged around 200 million years ago (Delsuc et al., 2018), these two species develop in a remarkably similar manner, with small cell numbers and shared stereotypic invariant cell lineages. Exactly 10 endoderm precursor cells actively drive endoderm invagination at the onset of gastrulation. This precise cellular framework is ideal to characterise the chain of molecular events occurring in each precursor, which collectively drive invagination (reviewed in Lemaire, 2011). Previous work established that invagination is a two-step process, conserved between *Phallusia mammillata* and *Ciona intestinalis* (Sherrard et al., 2010; Figure 1A). During step 1, endoderm cells apically constrict, leading to a flattening of the vegetal side of the embryo. During step 2, endoderm invagination proper takes place when endoderm cells shorten along their apico-basal axis while maintaining small apices. Both steps are controlled by Myosin II, via the phosphorylation of its regulatory light chain either at Ser19 (1P-Myosin) or at Ser19 and Thr18 (2P-Myosin) (Sherrard et al., 2010). During step 1, endoderm apical constriction is mediated by the apical accumulation of 1P-Myosin in response to Rho-associated kinase (ROCK). During step 2, sub-apical 2P-Myosin accumulation ensures that apical surfaces remain small, while baso-lateral 1P-myosin drives apico-basal shortening. Sub-apical 2P-Myosin accumulation is ROCK-dependent, but neither ROCK, RhoA, Rac nor Cdc42 are required for the baso-lateral accumulation of 1P-myosin. The two steps appear relatively independent of each other, as inhibition of step 1 does not prevent step 2 apico-basal shortening (Sherrard et al., 2010).

**Figure 1.**
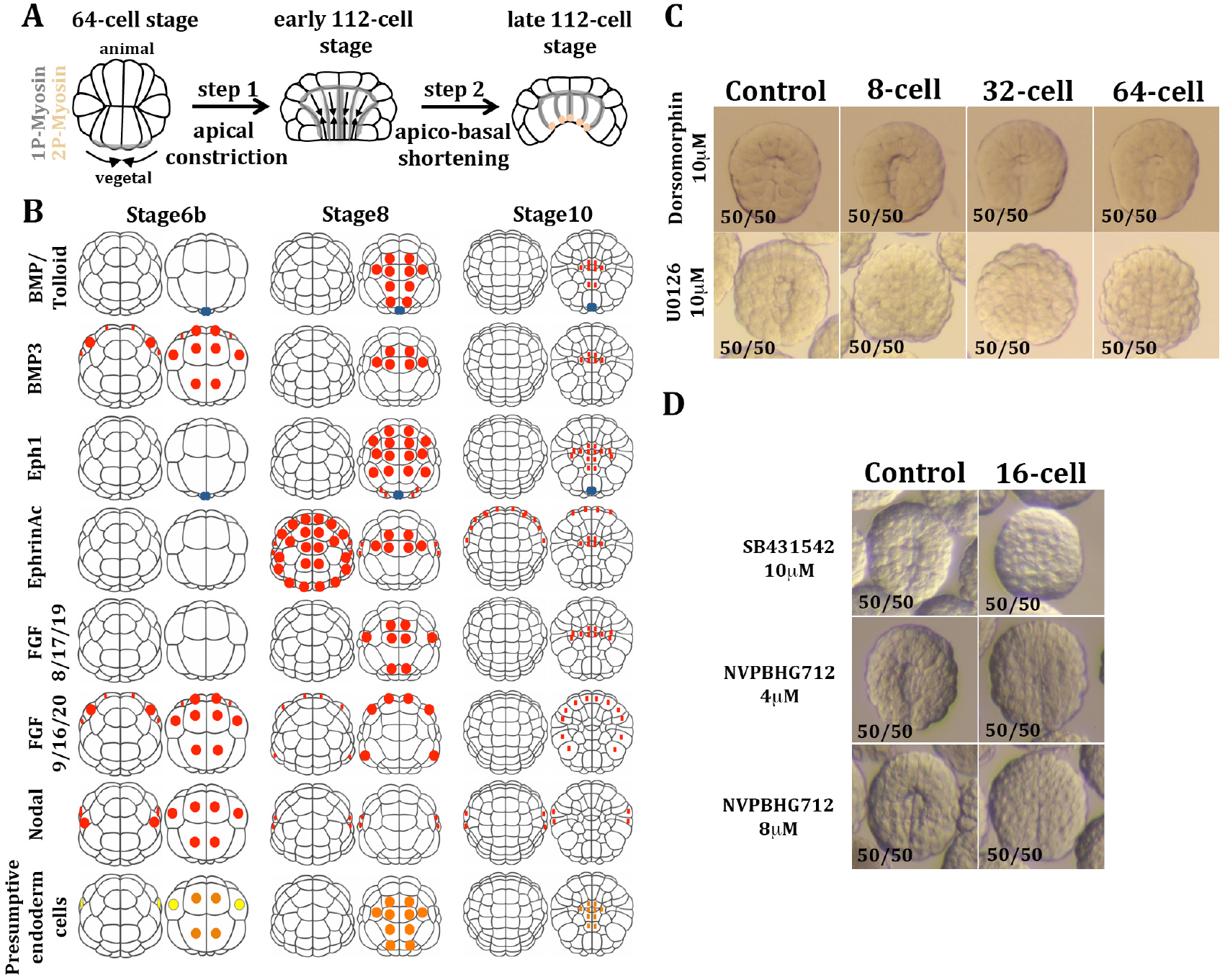
Nodal and Eph signalling regulate endoderm invagination in *Phallusia mammillata*. (A) Schematic representation of endodermal invagination in ascidian early embryos. (B) Schematic representations of the animal (left panels for each stage) and vegetal (right panels) hemispheres of *Ciona* embryos showing the expression pattern of developmental signalling ligands with predominantly vegetal expression in pre-gastrula stages (Imai *et al*, 2004; Yasuo *et al*, 2007; Hudson *et al*, 2005). Zygotic expression in shown in red, maternal mRNA in blue, fate-restricted primary endoderm precursors are in orange and pluripotent precursors giving rise to some primary endoderm are in yellow. At developmental stages 6b, 8 and 10 embryos have 32, 64 and 112 cells, respectively. (C) Late 112-cell stage *Phallusia mammillata* embryos treated with pharmacological inhibitors of BMP (Dorsomorphin at 10µM) and FGF (U0126 at 10µM; MEK inhibitor) signalling. Treatment was initiated at the 8-, 32- and 64-cell stages as indicated. (D) Late 112-stage *Phallusia mammillata* embryos treated with pharmacological inhibitors of Nodal (SB431542; Alk4/5/7 inhibitor) and Eph (NVPBHG712; Eph kinase inhibitor) receptor function from the 16-cell stage. In C and D, the inhibitors were dissolved in DMSO and control embryos were cultured in the same DMSO concentration as the inhibitor-treated embryos.

Several key questions remain unanswered. First, we do not know the identity of the apical activators of ROCK during step 1 and 2 or the pathway(s) leading to the baso-lateral accumulation of 1P-myosin during step 2. Second, what triggers the transition between the two steps remains mysterious. Third, the mechanisms that ensure that cell division is delayed in the endodermal lineage until after the completion of step 2 are still ill defined. Finally, the extent of coupling between the acquisition of the endodermal fate and the control of cell shape is also unknown. In this study, we addressed several of these issues. The results we report indicate that the apparently simple cell behaviours driving endoderm invagination are controlled by the sophisticated integration of multiple signalling pathways and can be partially uncoupled from the control of the endodermal cell fate.

## Results

### Nodal and Eph1 signals are required for endoderm invagination independently of mesendoderm specification

In *Drosophila* (Costa et al., 1994), *C. elegans* (Lee et al., 2006) or vertebrates (Heisenberg and Solnica-Krezel, 2008), cell behaviours during gastrulation are controlled by cell-cell communication, in part independently of cell fate specification processes in some of these systems. In *Ciona*, previous work revealed that ligands for the FGF, Wnt, Bmp, Nodal and Ephrin developmental signalling pathways are transcriptionally active in pre-gastrula endoderm precursors (Hudson et al., 2003; Imai et al., 2006, 2004; Shi and Levine, 2008; Yasuo and Hudson, 2007) (Figures 1B, S1). Early regulatory gene expression profiles in the vegetal territories of *Ciona* and *Phallusia* embryos are well conserved (Madgwick et al., 2018). We hypothesized that some of these signalling pathways may directly affect gastrulation.

Consistent with previous work (Hudson et al., 2003), inhibition of the FGF/ERK signalling with the MEK inhibitor U0126 (10 µM) from the 8- to 64-cell stages blocked invagination (Figure 1C). This invagination defect likely results from the fate switch of A-line endoderm precursors to trunk lateral cells (Shi and Levine, 2008), which do not invaginate at the onset of gastrulation. It was thus not investigated further. Similarly, while inhibition of the Wnt/ß-catenin pathway prevents gastrulation, this phenotype probably originates from mesendoderm fate misspecification (Hudson et al., 2013a; Imai et al., 2000), and was not studied in more detail. Mutation in *Ciona* of the non-canonical Wnt PCP pathway core component *Prickle*, transcriptionally activated from the 64-cell stage (Brozovic et al., 2018), does not affect gastrulation (Jiang et al., 2005), suggesting that unlike in vertebrates (Heisenberg and Solnica-Krezel, 2008), the Wnt PCP pathway is not required for the control of ascidian gastrulation.

Inhibition of Bmp signalling by overexpression of *Chordin* or *Noggin* in the distantly-related ascidian, *Halocynthia roretzi*, had no major effect on gastrulation, but prevented the formation of sensory head pigment cells (Darras and Nishida, 2001). Consistently, treatment of *Phallusia* embryos from the 8-, 32- or 64-cell stages with 10 µM of the Bmp receptor inhibitor Dorsomorphin (Yu et al., 2008) prevented the formation of the otolith pigmented cells without affecting or producing a delay in invagination, with the latter embryos eventually recovering and producing a nearly normal larva (Figure 1C, S3A and B, and not shown). Bmp is thus unlikely to play a major role during invagination.

Inhibition of the Nodal pathway was previously shown to impair gastrulation in *Ciona intestinalis* (Hudson et al., 2007) and *Nodal* expression is perfectly conserved between *Ciona* and *Phallusia* (Madgwick et al., 2018; Figure S2). Inhibition of Nodal signalling in *Phallusia mammillata* embryos by treatment from the 16-cell stage with SB505124 (50µM) or SB431542 (5-10µM), two selective pharmacological inhibitors of the Nodal receptor ALK4/5/7, blocked invagination (Figure 1D, 5, S4A), a phenotype also observed following the overexpression of a dominant negative form of the Nodal receptor ALK4/5/7 (*Ciinte.tAlk4/5/7*; Hudson *et al,* 2005) in *Phallusia* (Figure S4B). Interestingly, while in vertebrates Nodal controls both morphogenesis and mesendoderm fate specification (reviewed in Kiecker et al., 2016), inhibition of this signalling pathway in ascidian vegetal territories did not alter the specification of germ layers: expression levels of both early (Figure 2A) and late (Figure 2B) vegetal endodermal and mesoderm markers were not reduced by Nodal inhibition.

**Figure 2.**
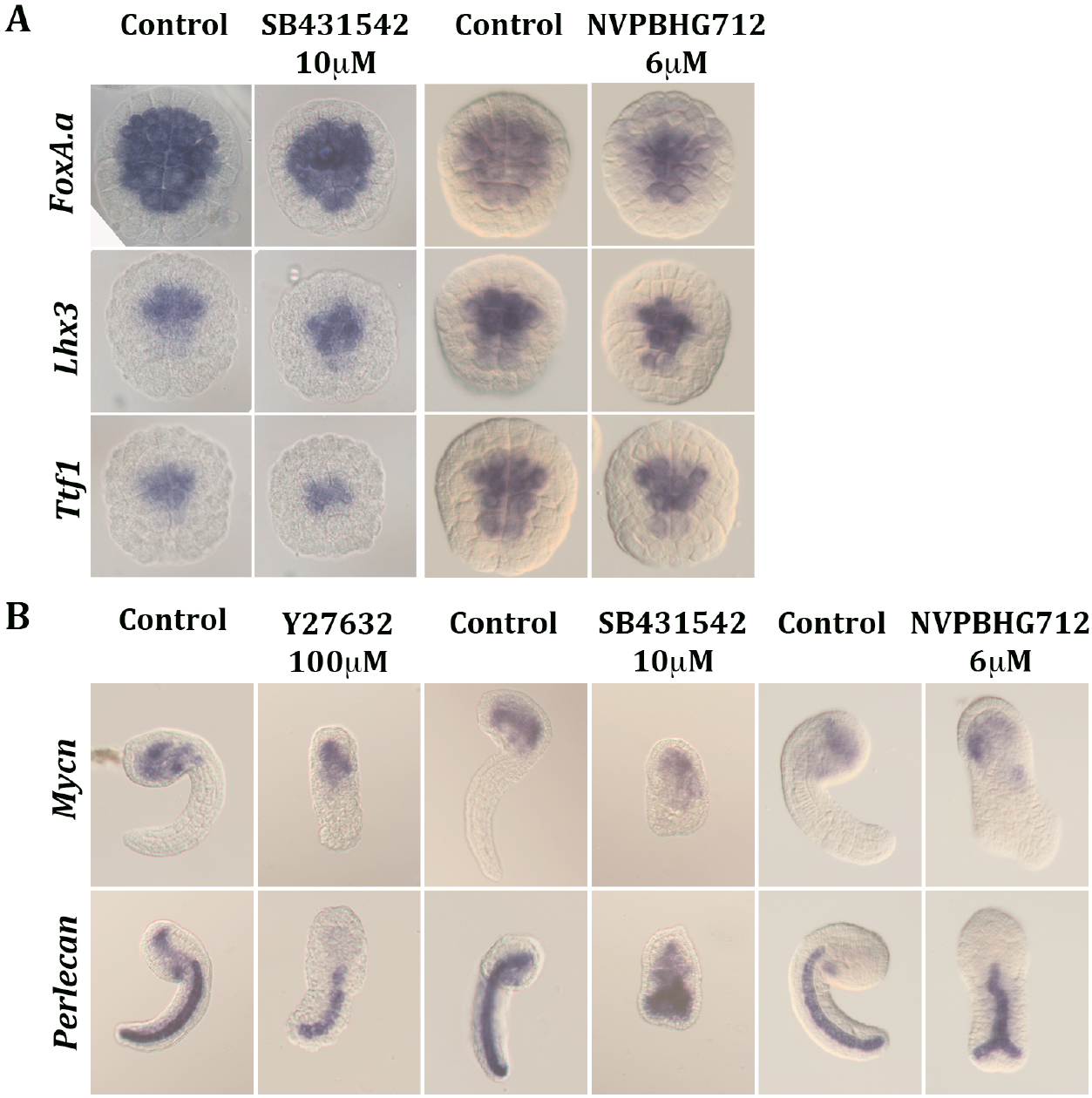
Nodal and Ephrin signalling are not necessary for vegetal mesendodermal cell fate specification. (A) Whole mount in situ hybridizations (WMISH) of *Phallusia* embryos at the early 112-cell stage using probes for *FoxA.a* (early mesendoderm marker), *Lhx3* and *Ttf1*, three transcription factors involved in endoderm cell fate specification. Control embryos were treated with DMSO. Nodal signalling was inhibited by treating the embryos with 10 µM SB431542 from the 16-cell stage. Eph mediated signalling was inhibited with 6µM NVPBHG712 from the 8-cell stage. (B) WMISH of *Phallusia* embryos at the mid tailbud II stage, using probes for *Mycn* (late endoderm marker) and *Perlecan* (late notochord marker) under control untreated conditions or following blockade of step 1 of endoderm invagination (100 µM of the ROCK inhibitor Y27632 from 64-cell stage) or of step 2 of endoderm invagination by inhibiting Nodal signalling (10 µM SB431542 from the 16-cell stage) or Eph signalling (6µM NVPBHG712 from the 8-cell stage). The inhibitors were dissolved in DMSO (SB431542, NVPBHG712) or H_2_O (Y27632) and control embryos were cultured in the same solvent concentration as the inhibitor-treated embryos.

We finally assessed the role of Eph/Ephrin signalling during invagination. Eph receptors having undergone independent gene duplication events in ascidians and vertebrates, each of the ascidian Eph receptors (the *Ciona* and *Phallusia* genomes harbour 6 and 5 Eph receptors, respectively) is orthologous to both EphA and EphB vertebrate receptors. Ascidian Eph3 receptor function is involved in anterior neural (Ohta and Satou, 2013), TLC (Shi and Levine, 2008) or nerve cord/notochord (Picco et al., 2007) inductions. Consistently, treatment of 16-cell stage *Ciona* embryos with 4-8µM NVPBHG712, a selective inhibitor of EphB kinases in mammalian systems (Martiny-Baron et al., 2010), phenocopied the effect of Eph3 signalling inhibition in early neural and notochord induction (Figure S5). In addition, it blocked endoderm invagination both in *Ciona* and *Phallusia* (Figure 1D, S4E) without preventing early or late vegetal endoderm or mesoderm marker gene expression (Figures 2, S6). Eph1 is the only Eph receptor specifically expressed throughout endoderm precursors at the time of endoderm invagination in *Ciona* and its expression profile is conserved in *Phallusia* (Madgwick et al., 2018; Figure S2). Consistent with the proposal that Eph1 is involved in endoderm invagination, this process was blocked by morpholino-mediated knockdown of *Eph1* in *Ciona* embryos (Figure S4 C-E).

We conclude from this section that endoderm invagination at the onset of ascidian gastrulation is independent of the Bmp and Wnt PCP pathways, controlled by FGF and Wnt/ß-catenin, probably indirectly through mesendoderm fate specification, and by Nodal and Eph1 signalling, independently of fate specification processes. Similarly, inhibition of Rok, the main driver of the first step of invagination, also did not affect endoderm and mesoderm specification, arguing for a broad uncoupling of morphogenetic and fate specification mechanisms. In the following sections, we further studied the mode of action and epistatic relationships of the Nodal and Eph pathways.

### Both Nodal and Eph1 signalling are required for the transition to step 2

As all ten endoderm precursors undergo simultaneous and similar shape changes during endoderm invagination (Sherrard et al., 2010), we focused our analyses on the cellular shape changes undergone by the A7.1 endoderm precursor. To understand which step of the invagination process was affected by Nodal and Eph signalling inhibition, we compared the height, apical and basal areas of the A7.1 cell in control embryos and in embryos treated with Nodal or Eph inhibitors or microinjected with Eph1 morpholinos (Figure 3 and S4 C-E). The invagination phenotypes described below were not due to a delay in development, as they persisted at later stages (data not shown).

**Figure 3.**
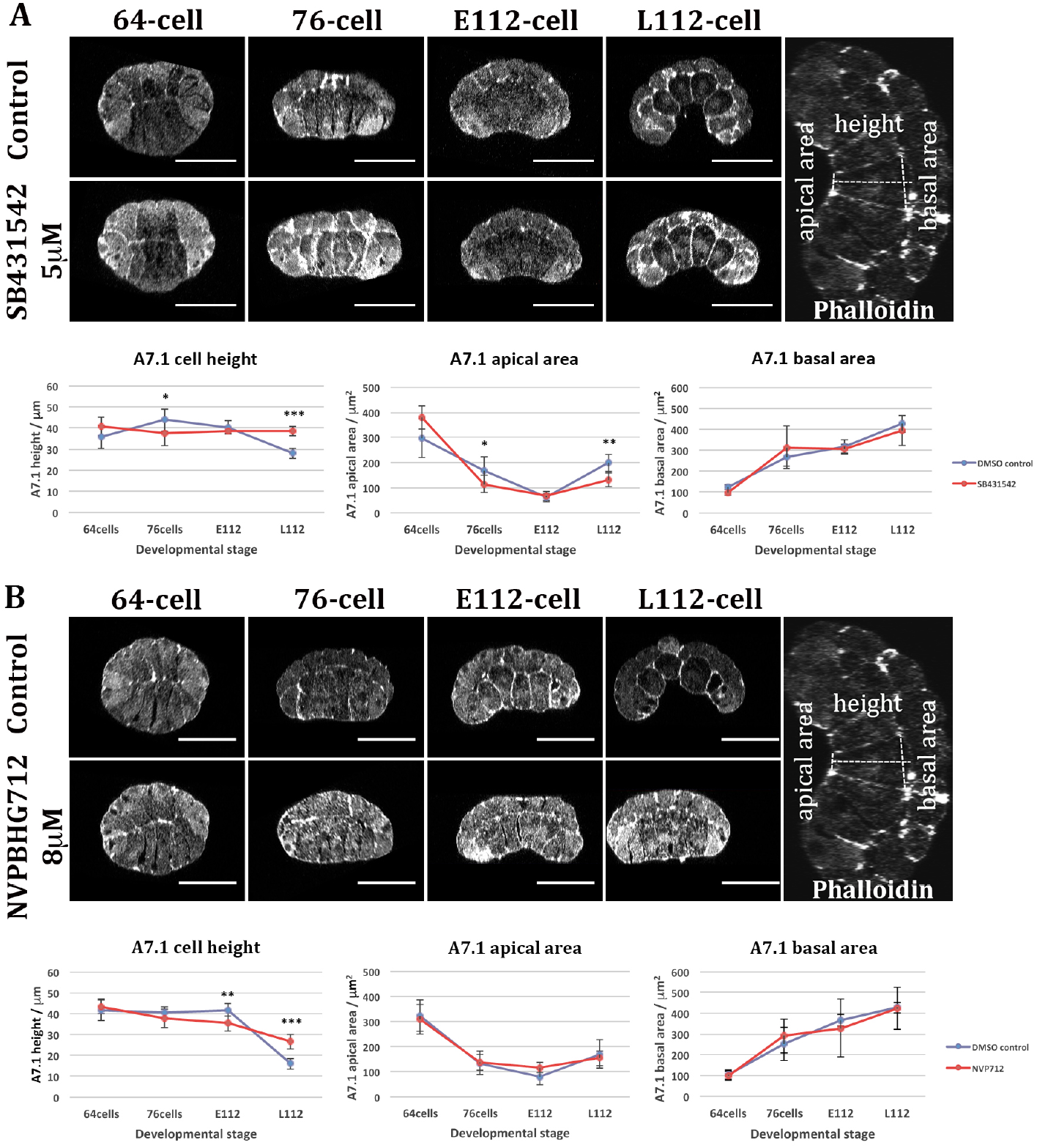
Nodal or Eph signalling inhibition causes defects in endoderm apico-basal shortening. (A) Frontal optical sections through *Phallusia mammillata* embryos stained with fluorescent Phalloidin to reveal cortical actin at different stages of endoderm invagination: 64-, 76-, early 112- (E112) and late 112-cell (L112) stages. The embryos were treated with either DMSO (Control) or with 5 µM SB431542 from the 16-cell stage. The bottom panels show a quantification of the evolution of the shape of the left endoderm A7.1 precursor in imaged embryos. Sample size: 4 < n < 12. () p>0.05, (*) p < 0.05, (**) - p < 0.01, (***) - p < 0.001 (t-test). (B) Frontal optical sections through *Phallusia mammillata* embryos stained with fluorescent Phalloidin to reveal cortical actin at different stages of endoderm invagination: 64-, 76-, early 112- (E112) and late 112-cell (L112) stages. The embryos were treated with either DMSO (Control) or with 8 µM NVPBHG712 from the 8-cell stage. Bottom panels: Quantification of the left A7.1 cell height, apical area and basal area of the embryos. M112-cell: mid 112-cell stage. Endoderm cells showing signs of premature cell division were not included in the quantitative analysis of endoderm cell shape dynamics. Sample size: 4<n<11. () p>0.05, (*) - p < 0.05, (**) - p < 0.01, (***)-p <0.001 (t-test). Scale bar: 50µm. Animal side is up and vegetal side is down. Scale bar - 50 µm.

Nodal receptor inhibition with SB431542 from the 16-cell stage in *Phallusia* embryos (Figure 3A) had no significant effect on the geometry of A7.1 up to the end of step 1 of invagination at the early 112-cell stage, and did not prevent apical constriction. By the late 112-cell stage, however, A7.1 cells were significantly taller in SB431542-treated embryos than in controls. Nodal signalling is thus dispensable for step 1 apical constriction but required for step 2 apico-basal shortening of endodermal precursors. Likewise, inhibition of Eph signalling by NVPBHG712 treatment from the 8-cell stage in *Phallusia* and *Ciona* embryos or following Eph1-MO injection in *Ciona* eggs left step 1 unaffected, while preventing correct apico-basal shortening of endodermal precursors during step 2 (Figure 3B and S4 C, D and E).

Localised myosin II contractility is the major driving force of ascidian endoderm invagination and is regulated by the phosphorylation state of its regulatory subunit (Sherrard *et al*, 2010). In control embryos during step 1 (apical constriction), 1P-myosin first accumulates in speckles on the apical surface of vegetal cells (Figure 4A, B control 76-cell stage), a signal which subsequently gradually decreases. During step 2 (apico-basal shortening), 1P-Myosin has disappeared from the apical side of endoderm progenitors and is detected on the baso-lateral surfaces of endoderm cells (Figure 4A, B control E112-and L112-cell stage).

**Figure 4.**
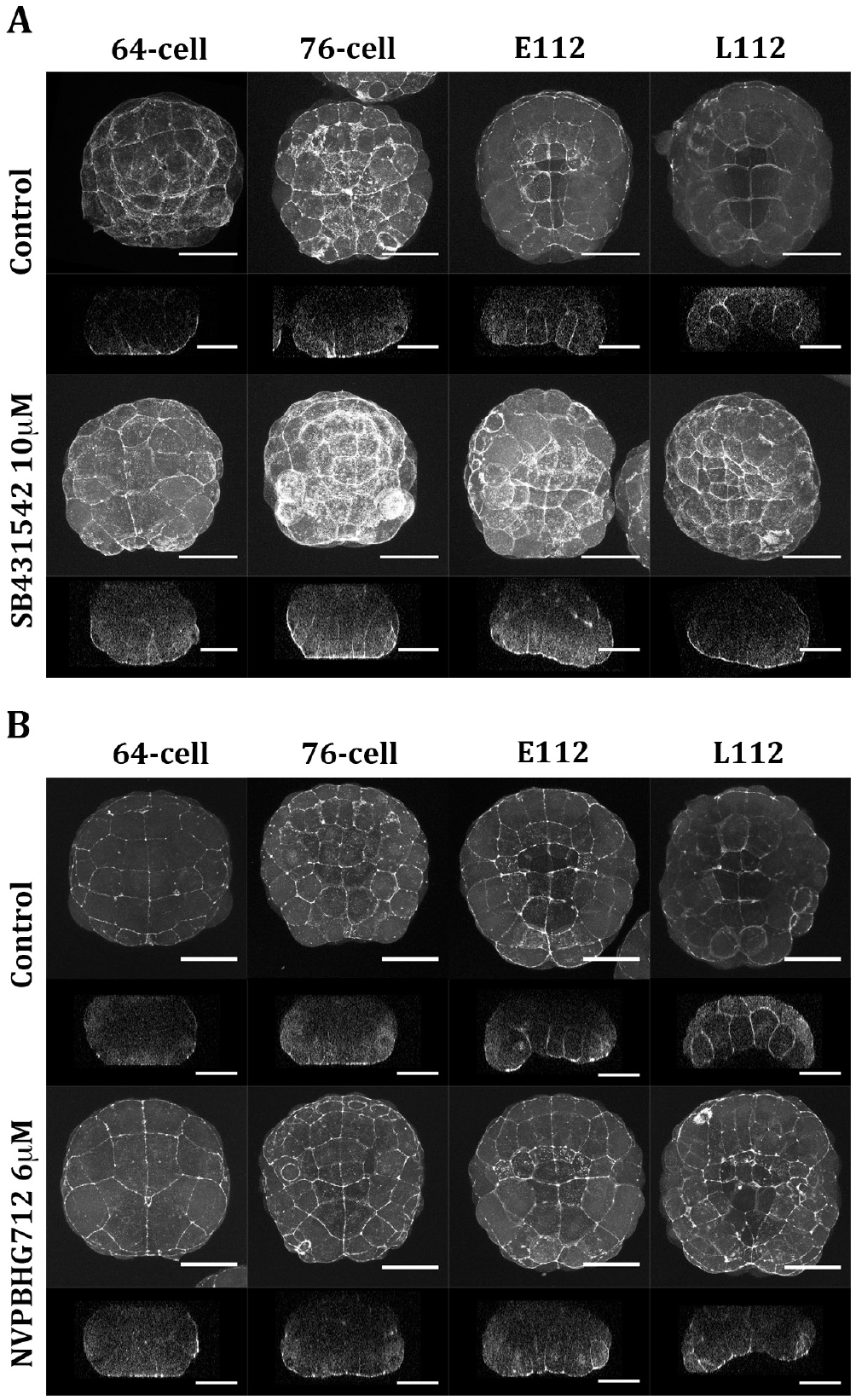
Nodal and Eph signalling regulate the transition between apical constriction and apico-basal shortening during endoderm invagination by modulating the pattern of 1P-Myosin. (A) Vegetal view of maximal projection (top) and frontal optical sections (bottom) of 1P-Myosin immunostainings of control (DMSO-treated) or Nodal receptor inhibited (10 µM SB431542 from 16-cell stage) *Phallusia mammillata* embryos fixed at the 64-cell, 76-cell, early 112-cell (E112) and late 112-cell (L112) stages. Scale bar: 50µm. (B) Vegetal view of maximal projection and frontal optical sections of 1P-Myosin immunostainings of control (DMSO-treated) or Eph receptor inhibited (6 µM NVPBHG712 from 16-cell stage) *Phallusia mammillata* embryos fixed at the 64-cell, 76-cell, early 112-cell (E112) and late 112-cell (L112) stages. 33% (2/6) of the treated embryos exhibited 1P-Myosin baso-lateral localisation, and successful invagination while all (6/6) embryos in the control conditions invaginated. In a second independent experiment, only 11% (1/9) of embryos treated with 6 µM NVPBHG712 from 16-cell stage invaginated and presented basolateral 1P-myosin accumulation, while 66.7% (6/9) control embryos invaginated. Scale bar: 50µm.

Upon Nodal signalling inhibition from 16-cell stage (Figure 4A, SB431542), the apical 1P-Myosin pattern of step 1 was established normally and possibly reinforced at 76-cell stage. 1P-Myosin apical accumulation, however, persisted throughout the 112-cell stage, while 1P-Myosin did not accumulate on the baso-lateral sides. Similarly, in Eph-inhibited embryos (Figure 4B, NVPBHG712), 1P-Myosin accumulated apically during step 1 as in wild-type but retained its apical localisation throughout the 112-cell stage without detectable baso-lateral reinforcement.

We conclude that Eph and Nodal signalling are required for the transition from step 1 to step 2. When either of these pathways is inhibited, step 1 occurs normally up to the early 112-cell stage. During the 112-cell stage, however, endodermal cells retain a step 1 pattern of 1P-Myosin localisation and never transition to step2.

### Nodal signalling sets the level of expression of *Eph1* in vegetal territories

We next explored the relationships between Nodal and Eph signalling during invagination. In both species, *Nodal* is first transiently expressed at the 32-cell stage in most vegetal cells, including the endoderm precursors (Figure S2, Imai et al., 2004). Its expression then becomes restricted to animal b-line ectodermal cells (Figure S2). SB431542 has been shown to rapidly penetrate cells and abrogate Nodal target gene expression within 40 min (Hudson et al., 2007). Early inhibition of Nodal signalling in *Phallusia* embryos either from the 16-cell stage onwards or for a shorter period between the 16-and 64-cell stages prevented endoderm invagination. Inhibition from the 64-cell stage onwards, however, had no major effect on endoderm invagination (Figure 5A). Nodal signalling during the 32-cell stage, when Nodal is expressed transiently in most vegetal cells, thus appears critical for endoderm invagination.

**Figure 5.**
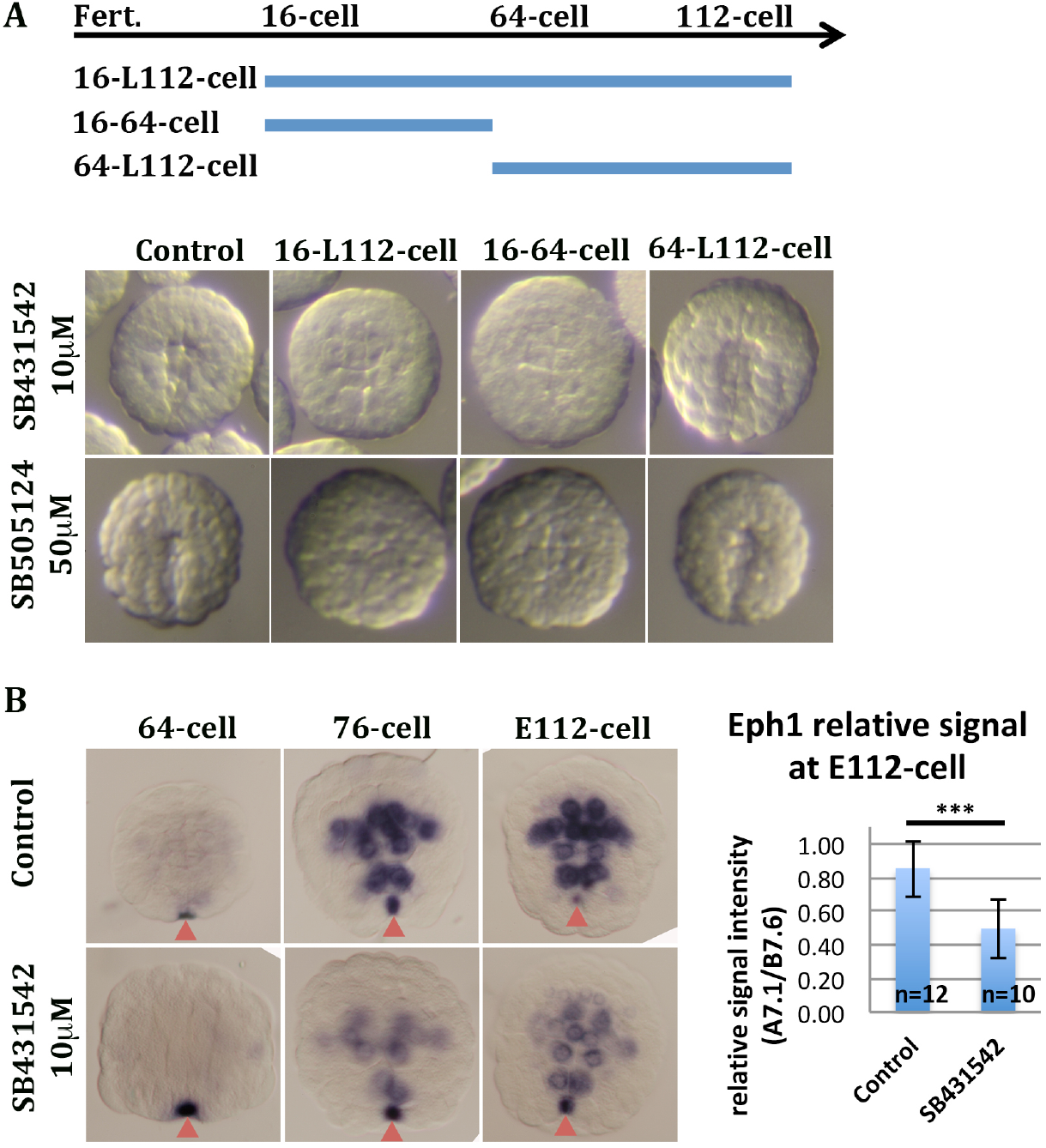
Vegetal Nodal signalling activity at the 32-cell stage regulates endoderm invagination and modulates *Eph1* expression in *Phallusia mammillata*. (A) Effects at the late 112-cell stage of treating embryos with Nodal signalling inhibitors (10µM SB431542 or 50 µM SB505124) during different developmental periods (top schema): 16-cell to late 112-cell stage (16-L112-cell stage), 16-cell to 64-cell stage (16-64-cell stage) and 64-cell to late 112-cell stage (64-L112-cell stage). (B) *Eph1* WMISH at the 112-cell stage in control embryos and in embryos treated with 10µM SB431542 from the 16-cell stage. Data representative of 3 independent experiments. Quantification of the relative *Eph1 in situ* signal at 112-cells stage with: *Eph1 relative signal intensity = (mean A7.1 signal – mean background signal) / (mean B7.6 signal – mean background signal)*. () p>0.05, (*) p < 0.05, (**) - p < 0.01, (***) - p < 0.001 (t-test). Note that SB431542 treatment decreases zygotic *Eph1* expression without affecting its localized maternal expression in the presumptive germ cells (red arrowheads), which formed a convenient internal control. Control embryos in A and B were incubated in DMSO at the same concentration as inhibitor-treated embryos.

This early requirement for Nodal signalling, more than an hour before the onset of invagination, suggests that this signalling pathway may indirectly regulate the transition to step 2 indirectly, via a transcriptional relay. *Eph1* is zygotically expressed from the 64-cell stage, approximately 45 minutes after the onset of *Nodal* expression in endoderm precursors (Figure S2, Imai et al., 2004). Vegetal *Eph1* expression, is initially very low, increases during endoderm invagination. To test whether Nodal signalling is regulating zygotic *Eph1* expression, *Phallusia* embryos were treated with SB431542 from the 16-cell stage and *Eph1* expression was assessed by whole mount *in situ* hybridisation (WMISH) (Figure 5B). Nodal signalling abolition did not qualitatively change the spatial domain of zygotic *Eph1* expression, indicating the presence of Nodal-independent *Eph1* transcriptional activators in endodermal precursors. Nodal signalling inhibition, however, markedly reduced the intensity of zygotic *Eph1 in situ* signal (Figure 5B), the intensity of the maternal signal (Fig 5B, arrowheads) serving as a reference.

We conclude that Nodal signalling sets the level, but not the spatial pattern, of expression of *Eph1* in the endoderm. The similarity of the phenotypes resulting from Nodal or Eph1 signalling inhibition suggests that Nodal regulates the transition to step 2 through the transcriptional upregulation of *Eph1* in endoderm precursors.

### Nodal-independent levels of Eph signalling are required to lengthen the cell cycle of endoderm precursors

We next analysed whether Nodal-independent Eph1 vegetal expression may also have a function during gastrulation. Cytokinesis and interphase cellular morphogenetic control need to be coordinated as they involve overlapping sets of cytoskeletal proteins (Duncan and Su, 2004). Gastrulating cells thus usually have a longer cell cycle than their neighbours, so that their mitosis is postponed until after completion of invagination. Indeed, cell division of ascidian endodermal precursors occurs after the completion of step 2 (Sherrard et al., 2010).

Full Nodal receptor inhibition or inhibition of Eph1 signalling with 6µM NVPBHG712 from the 16-cell stage, prevented step 2 without interfering with the timing of division of endoderm progenitors (not shown). By contrast, treatment of embryos with a higher concentration (8µM) of NVPBHG712 led to a premature division of endoderm progenitors in 15.6% of *Phallusia* treated embryos (Figure S8A). Similarly, *Eph1* morpholino injection in *Ciona* led to the premature division of endoderm precursors (Figure S8B).

We conclude that Eph1 has a dual role during endoderm invagination. Nodal-independent low levels of Eph1 expression are required to lengthen the cell cycle of endodermal precursors and postpone cell division until the completion of step 2. Nodal-dependent higher levels of Eph1 expression are then required for the transition between steps 1 and 2.

## Discussion

We studied the regulatory logic of the invagination of endodermal progenitors at the onset of ascidian gastrulation. This invagination involves a myosin-dependent two-step change of shape of invaginating cells and a lengthening of their cell cycle. We identified two novel regulators of these processes: the Nodal TGF-ß pathway and the Eph1 receptor tyrosine kinase pathway (see model on Figure 6). Nodal, acting via the transcriptional upregulation of *Eph1*, is necessary for the transition between the two steps of cell shape changes. In addition, low, Nodal-independent, Eph1 signalling is required for the lengthening of the endodermal cell cycle. Interestingly inhibition of Nodal or Eph signalling had no major effect on the fate of mesendodermal cells, demonstrating a partial uncoupling between fate specification and morphogenetic processes in ascidian embryos.

**Figure 6.**
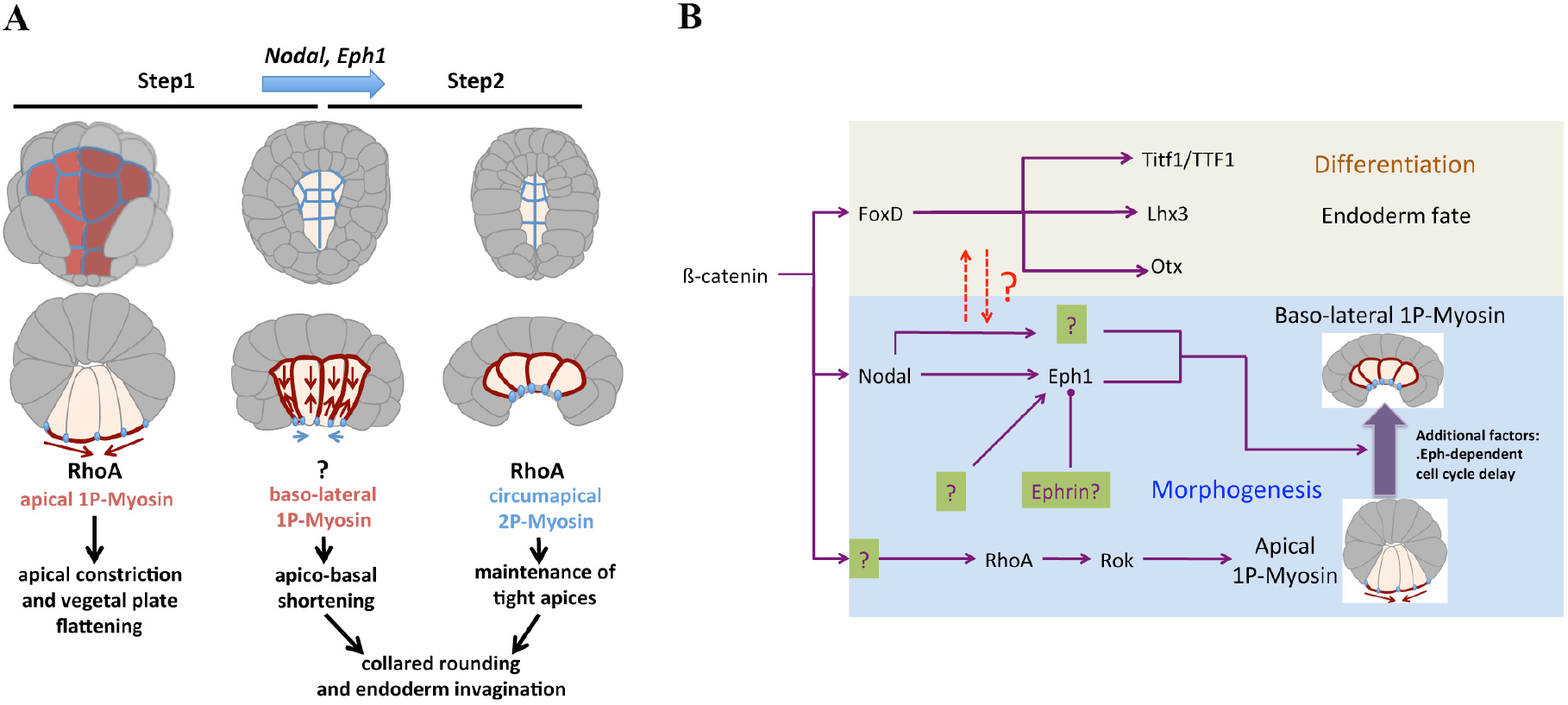
The emerging endoderm invagination regulatory network. (A) Regulatory mechanisms involved in endoderm cell shape changes during the 2-step process of ascidian endoderm invagination. (B) Emerging transcriptional regulatory network driving endoderm invagination. The scheme highlights the unknown relationship between cell fate regulation and cellular morphogenesis.

### An active transcriptional switch between apical constriction and apico-basal shortening

We previously showed that invagination involved a first step of apical constriction, driven by the activation of Myosin II by Rho kinase on the apical surface of endodermal progenitors, followed by a second step of apico-basal shortening driven by the Rho kinase-independent activation of myosin II on the baso-lateral facets of invaginating cells (Sherrard et al., 2010). These two steps appear to be independent of one another as inhibition of apical constriction does not affect apico-basal shortening (Sherrard et al., 2010). A similar situation is found during *Drosophila* gastrulation, as apical constriction can be blocked without affecting subsequent apico-basal shortening (Leptin, 1999). More generally, cells frequently switch from one behaviour to another during animal gastrulation (Davidson, 2012; Leptin, 2005). How the transitions between successive and independent modules are controlled at the molecular level remains mysterious and could involve direct mechanosensory feedbacks or transcriptional processes.

By searching for signalling pathways involved in ascidian endoderm invagination, we discovered that Eph signalling inhibition stalls invaginating cells in a state corresponding to the end of apical constriction (Figure 4). This finding carries two major conceptual messages. First, it suggests that, in the absence of apico-basal shortening, the Rho kinase-driven apical constriction step is stabilized and prolonged. Second, it identifies the transcriptional upregulation of Eph as a key component of the switch between the two steps. While we cannot rule out the existence of direct mechanosensory feedback mechanisms within the cell cortex, these results combined with the normal onset of step 2 when step 1 is blocked (Sherrard et al., 2010), argue that the transition between the two steps is controlled at the transcriptional level by developmental gene regulatory networks.

The observed stalling of cells in an end-of-step-1 state could result from two different mechanisms. The simplest explanation is that Eph activation on the apical side of endodermal cells directly terminates 1P-myosin apical accumulation at the end of step 1. As myosin is activated by Rho kinase during step 1, this scenario would be compatible with the report that EphA4 kinase activity inhibits RhoA in early *Xenopus* embryos (Winning et al., 2002). Alternatively, Eph1 could be activated during step 2 on the basolateral side of cells, where it would activate myosin. In this scenario, the maintenance of apical 1P-myosin when Eph signalling is decreased suggests a negative feedback between basolateral and apical 1P-myosin accumulation. To discriminate between these two scenarios, it will be necessary to identify the intracellular effectors of high Eph1 signalling and their subcellular localization.

### Eph1 signalling independently controls cell cycle length in endodermal progenitors

As mitosis and myosin-driven cell shape changes during interphase use overlapping sets of cytoskeletal proteins, cell division cannot be executed during the invagination of epithelial cells (Duncan and Su, 2004). In pre-gastrula and early gastrula ascidian embryos, the mitosis of endodermal progenitors is thus delayed until the end of their invagination (Nishida, 1987). Our finding that Eph1 inhibition triggers precocious mitosis in these precursors constitutes the first identification of a zygotic control mechanism of mitosis timing in ascidians. As Eph1 inhibition does not alter the fate of endodermal cells (Figure 2), the timing of mitosis appears to be controlled by cell-cell communication rather than by a cell-autonomous fate-specific timer.

In *Drosophila* and *Xenopus,* mitotic delay is achieved through the control of the phosphorylation of CDK1 by the inhibitory CDC25 phosphatase (Grosshans and Wieschaus, 2000; Murakami et al., 2004; Seher and Leptin, 2000). Eph signalling has previously been implicated in both positive (Genander et al., 2009) and negative (del Valle et al., 2011) control of cell proliferation in mammals. Eph receptor activation can *in vivo* phosphorylate the Src kinase (Jungas et al., 2016), which is a negative regulator of CDK1 (Horiuchi et al., 2018). *In vitro*, many Eph receptors can also directly phosphorylate CDK1 on its inhibitory regulatory Tyr15 residue (Blouin et al., 2011). Future studies will explore whether Eph1 delays mitosis of endodermal progenitors via the inhibition of CDK1 activity in ascidians as well.

### Integration of morphogenetic and fate specification cues

*Nodal* is broadly expressed in most vegetal cells at the 32-cell stage, including cells that will not invaginate. Its morphogenetic activity thus needs to be gated by fate specification cues to only affect endoderm precursors. Our work indicates that this gating occurs at the level of *Eph1* expression. We indeed find that two independent regulatory inputs drive the expression profile (Figure 5) of Eph1. First, fate specification regulatory networks ensure that this gene is specifically expressed in endoderm precursors from the 64-cell stage (Figure S2). Second, Nodal-dependent morphogenetic cues ensure that *Eph1* reaches a sufficient level expression to trigger the invagination of endodermal cells.

An open question in this work is the identity of the ligand that elicits Eph1 signalling during endoderm invagination. Four closely related paralogous *Efna* genes, *Efna.a* to *Efna.d*, exist in ascidians, clustered on the same chromosome (Mellott and Burke, 2008). Of these, only *Efna.c* is expressed in endoderm precursors prior to gastrulation. We verified that Nodal signalling inhibition, has no effect on *Efna.c* expression (Figure S7). It remains unclear what is the relevant ligand for Eph1 activity regulating endoderm invagination.

### Relationships between fate specification and morphogenesis in chordates

Our work identifies Nodal-dependent actomyosin contractility as a major force during ascidian endoderm invagination. In vertebrates, Nodal also controls gastrulation movements (Heisenberg and Solnica-Krezel, 2008) and, in zebrafish at least, this action involves the control the actomyosin contractility of mesendodermal cells. The control of cortical tensile forces by Nodal thus appears to be a shared feature of ascidians and vertebrates. Interestingly, in *Xenopus*, Nodal signalling is necessary for normal EphA4 expression (Wills and Baker, 2015) in the mesoderm and both signalling pathways are required for the internalization of the mesendoderm (Evren et al., 2014). The presence of this association in both taxa could reflect either convergence or an ancestral state.

One crucial difference between ascidians and vertebrates is that in the latter, Nodal controls both mesendoderm fate specification and morphogenetic movements. In *Xenopus* (Luxardi et al., 2010) and zebrafish (Hagos and Dougan, 2007; Sun et al., 2006), the two activities can, however, be temporally uncoupled, the period of mesendoderm induction controlled by Nodal signalling ending before the onset of gastrulation. Furthermore, in *Xenopus*, distinct Nodal paralogues sequentially control mesendoderm fate specification and gastrulation movements (Luxardi et al., 2010). A partial uncoupling between the morphogenetic and patterning roles of Nodal is thus also observed in at least some vertebrates. Interestingly, while in ascidian embryos Nodal controls invagination, in *Amphioxus* embryos this pathway is required for antero-posterior patterning of the mesendoderm, but not for invagination (Onai et al., 2010). Thus, while Nodal likely plays an ancestral role at the time of gastrulation in all three chordate groups, its precise role in the chordate ancestor remains elusive.

Several scenarios could explain the evolution of the function of Nodal. First, mesendoderm fate commitment, which is completed prior to the onset of gastrulation in ascidians (Lemaire, 2009), occurs during invagination in vertebrates (Ho and Kimmel, 1993; Kato and Gurdon, 1993). Invaginating vertebrate cells thus still need to communicate with each other to acquire their proper fate, which may benefit from a tighter integration of cell fate and morphogenesis control mechanisms. Second, mesendoderm internalization involves a much larger palette of cell type-specific cellular behaviours in vertebrates than in ascidians or cephalochordates, including epithelial to mesenchymal transition, cell migration, ingression or convergence and extension. This more complex cellular organization might require fate and cell shape regulation to be more tightly coupled than in ascidians or cephalochordates where invaginating cells retain their epithelial organization. Consistent with the notion that a simpler epithelial invagination may more easily be controlled independently of fate specification mechanisms, *Drosophila melanogaster* mesoderm invagination is controlled the *folded gastrulation* signalling pathway, which controls the pattern of Myosin II localization without controlling cell fate specification (Costa et al., 1994; Dawes-Hoang et al., 2005; Leptin and Grunewald, 1990).

Simpler embryos thus appear to operate partially morphogenetic sub-regulatory networks orchestrating cell shape changes independently of cell fate specification during mesendoderm invagination. This offers the opportunity to study morphogenesis without the added complication of cell fate specification mechanisms.

## Materials and Methods

### Embryo culture conditions

Adult *Phallusia mammillata* and *Ciona intestinalis* (formerly known as *Ciona intestinalis* type B) (Brunetti et al., 2015; Pennati et al., 2015) were collected on the Northern shore of Britany by the Marine facility of the Roscoff Marine Biological Station (France) and maintained in natural or artificial sea water at 16°C under constant illumination. Eggs were collected, fertilized and dechorionated as previously described (McDougall and Sardet, 1995; Robin et al., 2011).

### Perturbation assays by chemical treatment

*Phallusia* embryos were treated with the MEK inhibitor U0126 from Calbiochem (10µM), the BMP signalling inhibitor Dorsomorphin from SIGMA (10µM), the Nodal receptor inhibitors SB431542 (5µM and 10µM) and SB505124 from SIGMA (50µM), the Eph inhibitor NVPBHG712 from Tocris Bioscience (1, 2, 4, 6 or 8 µM), and with the Rho kinase inhibitor Y-27632 from SIGMA (100µM) in artificial sea water at specific developmental stages as specified in the figures. *Ciona* embryos were treated with SB431542 and NVPBHG712 at 5 µM and 8 µM, respectively, at the developmental stages specified in the figures. SB431542 and SB505124 are reported to be selective potent inhibitors for Alk4, Alk5 and Alk7 TGFβ receptors (Inman et al., 2002; DaCosta Byfield et al., 2004) and therefore inhibit the sole ascidian orthologue of these receptors. NVPBHG712 has been described as a highly selective small molecular weight inhibitor of Eph kinase activity (Martiny-Baron et al., 2010). SB431542, U0126, Dorsomorphin and Y27632 are classically used inhibitors in ascidian developmental studies (Hudson et al., 2007, 2003; Sherrard et al., 2010; Waki et al., 2015).

### Gene identities

| Gene name | *Ciona intestinalis*<br>Unique gene identity | *Phallusia mamillata*<br>Unique gene identity |
| --- | --- | --- |
| *Alkaline phosphatase* | *Cirobu.g00011480* | *Phmamm.g00011476* |
| *Alk4/5/7* | *Cirobu.g00012156* | *Phmamm.g00004838* |
| *Beta-catenin* | *Cirobu.g00010084* | *Phmamm.g00012274* |
| *Bmp1* | *Cirobu.g00002684* | *Phmamm.g00000600* |
| *Bmp3* | *Cirobu.g00003050* | *Phmamm.g00001174* |
| *Brachyury* | *Cirobu.g00013860* | *Phmamm.g00007005* |
| *Eph1* | *Cirobu.g00000642* | *Phmamm.g00004451* |
| *Eph3* | *Cirobu.g00008427* | *Phmamm.g00005695* |
| *Ephna.c* | *Cirobu.g00005705* | *Phmamm.g00000939* |
| *Ephna.d* | *Cirobu.g00005918* | *Unclear identity* |
| *Etr* | *Cirobu.g00007645* | *Phmamm.g00007762* |
| *Fgf8/17/18* | *Cirobu.g00007390* | *Phmamm.g00011773* |
| *Fgf9/16/20* | *Cirobu.g00004295* | *Phmamm.g00003805* |
| *FoxA.a* | *Cirobu.g00002136* | *Phmamm.g00001891* |
| *FoxD* | *Cirobu.g00009025* | *Phmamm.g00006179* |
| *Lhx3* | *Cirobu.g00014215* | *Phmamm.g00016546* |
| *Mrf* | *Cirobu.g00003985* | *Phmamm.g00010708* |
| *Mycn* | *Cirobu.g00012221* | *Phmamm.g00007048* |
| *Nodal* | *Cirobu.g00010576* | *Phmamm.g00015500* |
| *Perlecan* | *Cirobu.g00005372* | *Phmamm.g00005761* |
| *Ttf1* | *Cirobu.g00001550* | *Phmamm.g00010419* |
| *Twist-like-1* | *Cirobu.g00007069* | *Phmamm.g00000523* |

### Microinjection of RNAs and morpholino oligonucleotides

Synthetic mRNA was produced using the mMESSAGE mMACHINE kit (Ambion) and microinjected as described in (Hudson et al., 2003). Synthetic mRNA for the *Ciona* dominant negative Nodal receptor *Alk4/5/7* (Hudson, 2005) was microinjected at a concentration of 1.190 µg/µl. Morpholino-antisense oligonucleotides against *Ciinte Eph1*, Eph1-MO (ATCTCCAATCTCCGGTCTGTTTGTC), were dissolved in water at a concentration of 0.7 mM before microinjection.

### Whole-mount *in situ* hybridization

*In situ* hybridization experiments in *Phallusia* embryos were performed as described in Christiaen *et al*, 2009. Dig-labelled mRNA probes for *Phallusia mammillata FoxA.a*, *Lhx3*, *Mycn*, *Perlecan* and *Ttf1* were synthesized from cDNA clones of the Villefranche-sur-mer *Phallusia mammillata* cDNA clone collection (Brozovic et al., 2018) using the SP6/T7 DIG RNA labelling kit (Roche, ref. 11175025910). Clones used for probe synthesis: *FoxA.a* – AHC0AAA74YF05, *Lxh3* – AHC0AAA183YA13, *Mycn* – AHC0AAA60YB16, *Perlecan* - AHC0AAA215YL21, *Ttf1*-AHC0AAA267YK08.

*Nodal* (phmamm.g00015500), *Ephna.c* (phmamm.g00000939) and *Eph1* (phmamm.g00004451) Dig-labeled mRNA probes were synthesized from cDNA produced from total mRNA (Superscript III Reverse transcriptase kit; Life Technologies) isolated from *Phallusia* embryos at 32-cell stage for *Nodal* and at 112-cell stage for *Ephna.c* and *Eph1* (RNeasy minikit; Qiagen). The primers introduce a T7 promoter (in bold) making the PCR product directly a probe synthesis template. The primers used to amplify the cDNA templates were:

*Nodal* (phmamm.g00015500)

Forward: 5’-CTATGGATATGACACAAGTATCGTTCTGC-3’;

Reverse: 5’-GATCC**TAATACGACTCACTATAGGG**TTATCGACATCCACATTCT-3’;

*Eph1* (phmamm.g00004451)

Forward: 5’-CCAACGTTGCGACTCCACTTTCACC-3’

Reverse: 5’-ACGA**TAATACGACTCACTATAGGG**GTCTTGGTTAGACATCTCCCAG-3’;

*Ephna.c* (phmamm.g00000939)

Forward: 5’-CAACGAGGCATGTTCTCTATTGGA3’;

Reverse: 5’-GGATCC**TAATACGACTCACTATAG**GATATGACGAAACCAGCAGTCAC-3’;

The protocols for *in situ* hybridization and alkaline phosphatase staining of *Ciona* embryos were described previously in Hudson *et al*., 2013. Probes used include *Brachyury*, *Etr*, *Lhx3*, *Mrf*, *Ttf1* and *Twist-like-1* as previously described (Corbo et al., 1997; Hudson et al., 2013a; Meedel et al., 2007; Ristoratore et al., 1999; Hudson and Yasuo, 2006; Hudson et al., 2003).

### Immunohistochemistry

Phalloidin-stained embryos were fixed in 4% formaldehyde in artificial sea water with HEPES (ASWH), at room temperature for 10 minutes when not imaged in Murray’s clear solution. Embryos imaged in Murray’s clear solution were fixed in fixation solution (4% formaldehyde, 50mM EGTA, 100 mM PIPES pH 6.9, 400 mM sucrose) at room temperature for 10 minutes. After fixation the embryos were washed 3 times in PBT (0.1% Tweens in PBS), once with PBS and stained with Phalloidin (5µl Phalloidin per ml of PBS; Alexa Fluor546 phalloidin; A22283 Life Technologies) at 4°C, overnight. The stained embryos were washed 3 times with PBS and either mounted directly with mounting media (80% glycerol, 10% 10XPBS, 1.6 % propyl gallate in H_2_O) or dehydrated by going through an isopropanol series (70%, 85%, 95%, 100%, 100%) and then cleared by washing 3 times in Murray’s Clear solution (benzyl benzoate:benzyl alcohol; 2:1) and imaging in Murray’s Clear solution.

Anti-phospho-myosin-stained embryos were fixed at room temperature for 30min in fixation solution (100mM HEPES pH7.0, 0.5mM EGTA, 10mM MgSO_4_, 300mM Dextrose, 0.2% Triton, 2% Formaldehyde EM-grade, 0.2% Gluteraldehyde). Embryos were then washed 3 times with PBT, once with PBS and quenched with 0.1% sodium borohydride in PBS for 20min. They were blocked with 1% BSA in PBT for 24h and stained with primary antibody recognizing Ser19 phospho-myosin (1:50; rabbit; 3671S Cell Signaling Technology) for 24h. The samples were washed 3 times in PBT and stained with a donkey anti-rabbit FITC or Alexa647 labelled secondary antibody (1:200; 711-095-152 Jackson ImmunoResearch or A21244 Molecular Probes), washed 3 times with PBT and mounted in mounting media (Sherrard et al., 2010).

### Imaging and image analysis

*Phallusia* confocal fluorescence imaging was performed with a Zeiss LSM780 using a 40X objective (NA 1.3), 1.2X zoom and a step of 1µm between sections. Imaged embryos were oriented with Amira and cell dimensions measured with Fiji (Schindelin et al., 2012). Apical and basal area measurements in *Phallusia* correspond to the area along the last plane of contact with lateral neighbour cells and not to the curved apical and basal surfaces. *In situ* Eph1 relative signal intensities were determined on inverted *in situ* images using Fiji. The *in situ* images had the dimensions of 5185 x 3456 pixels. For the relative Eph1 signal intensity calculation the mean signal intensity in a constant area was measured in cells A7.1 (endoderm) and B7.6 (germ line) with a single measurement (area measured: 7556 pixels), as well as the mean background signal intensity with the average of three independent measurements per image (area measured: 485200 pixels). The relative signal intensities were calculated using the formula: *Eph1 relative signal intensity = (A7.1 mean signal - background signal) / (B7.6 mean signal – background mean signal)*.

Confocal imaging of *Ciona* embryos was carried out with a Leica SP5 using a 40X objective (NA 1.25), 1.5X zoom and a z-step of 1µm. To measure apical surface areas and cell heights along the apico-basal axis of *Ciona* endoderm precursors, embryos were individually mounted and oriented so that the regions of interest face roughly to the objective. ImageJ (Schneider et al., 2012) was used to further orient acquired images and for measurements. For the *Ciona* images shown in Figure S4A, fixed embryos were placed with their vegetal pole side up and photographed with a Leica Z16 APO with a Canon EOS 60D mounted on it.

## Acknowledgments

We thank J. Piette (CRBM, Montpellier) for reagents and discussions. We thank F. Robin (UPMC, Paris) and C. Hudson (LBDV, Villefranche sur Mer) for technical advice and discussions. We thank the Montpellier Resource Imaging (MRI) facility, which provided the confocal Zeiss LSM780 microscope. Work in PL’s team was supported by the Centre National de Recherche Scientifique (CNRS) and the Agence Nationale de la Recherche (contracts Geneshape-ANR-SYSC-018-02 -, Chor-Evo-Net – ANR 2008 BLAN 0067 91 – and Dig-Em - ANR-14-CE11-0013). The team of H. Y. was supported by the Centre National de la Recherche Scientifique (CNRS), the Sorbonne Université, and the Agence Nationale de la Recherche (ANR-09-BLAN-0013-01)). U.M. Fiúza was funded by the GeneShape ANR contract followed by an FRM (SPF20120523969) postdoctoral fellowship.

## Author contributions

All *Phallusia mammillata* experiments were done by UM Fiuza. The assays testing the specificity of the NVPBHG712 pharmacological inhibitor in *Ciona intestinalis* were carried out by A. Rouan. All the remaining *Ciona* experiments were performed by T. Negishi and H. Yasuo. The manuscript was written by UM Fiuza, H. Yasuo and P. Lemaire with comments from all authors.

