## Supplementary Material for "A Nodal/Eph signalling relay drives the transition from apical constriction to apico-basal shortening in ascidian endoderm invagination"

### The supplementary material section includes 8 supplementary figures (Figs S1 to S8)

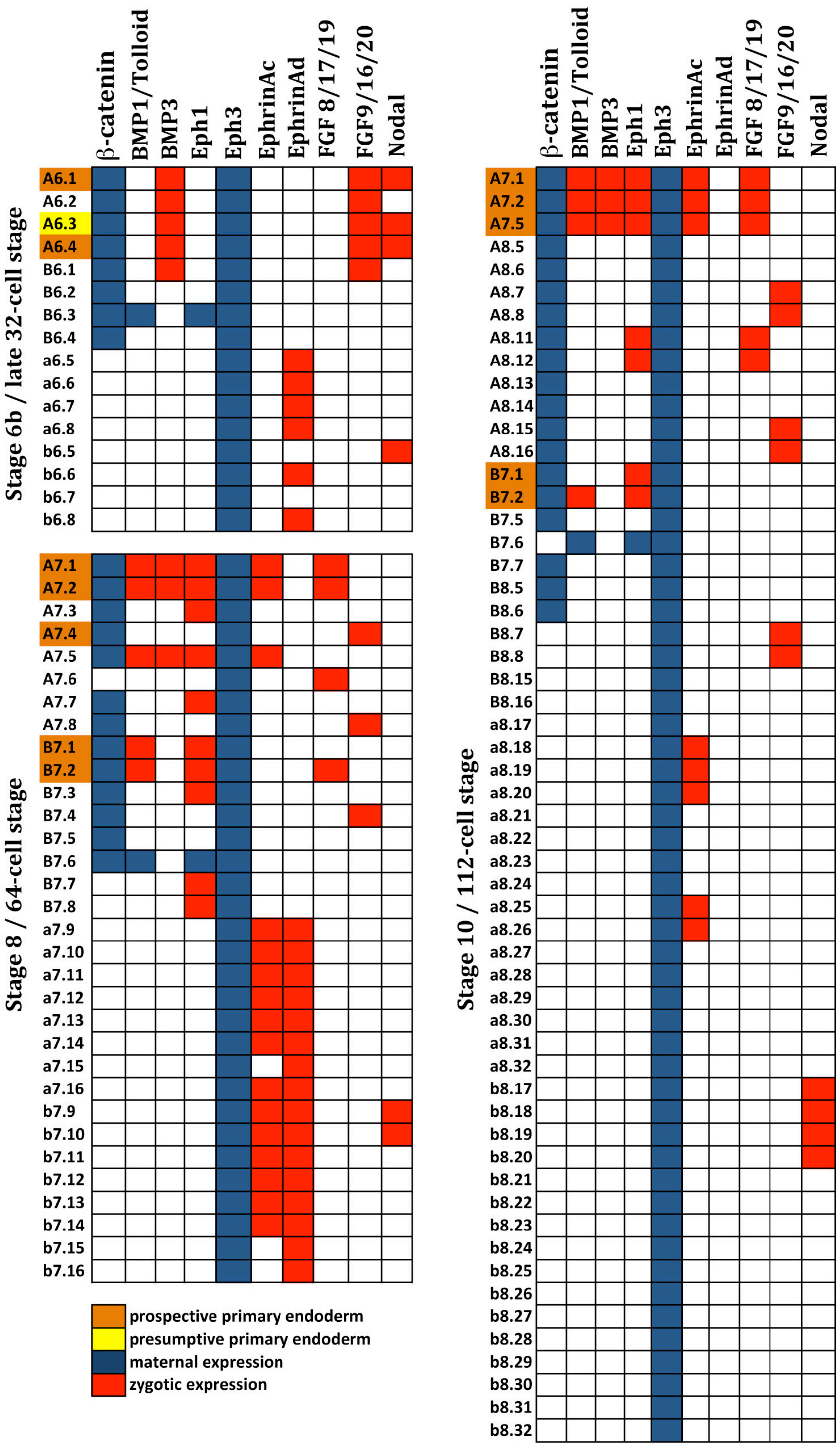


**Figure S1. Expression profile of *Ciona* signalling genes with predominant vegetal expression and related regulatory genes**

Expression pattern of developmental regulatory ligands with predominant vegetal expression in pre-gastrula stages at stage 6b, stage 8 and stage 10 (Imai *et al*, 2004; Yasuo *et al*, 2007; Hudson *et al*, 2005). Zygotic expression in red, maternal mRNA in blue, primordial endoderm cells in orange and presumptive endoderm cells in yellow.

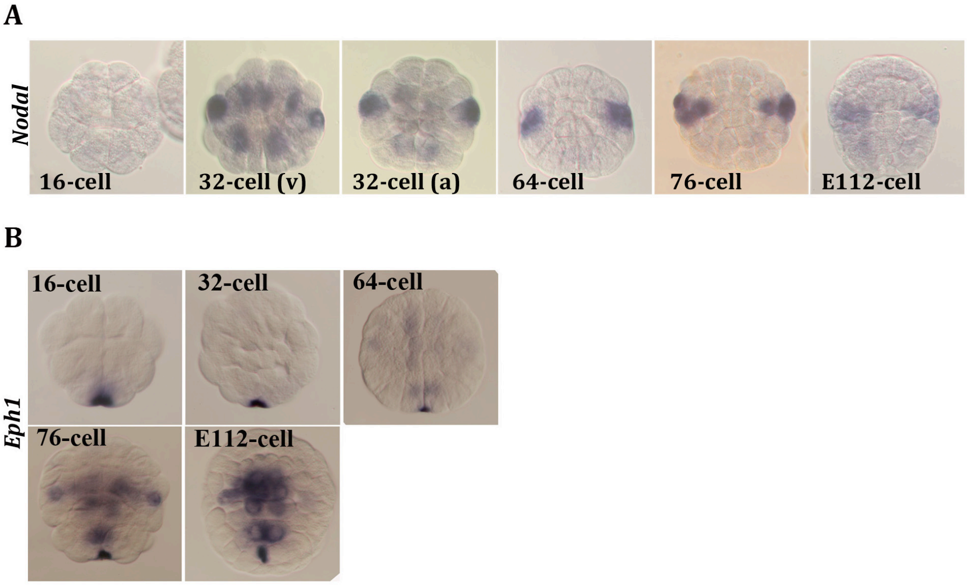


**Figure S2. *Nodal* and *Eph1* gene expression patterns are conserved between *Phallusia mammillata* and *Ciona intestinalis***

(A) Whole mount *in situ* hybridization (WMISH) of *Nodal* in *Phallusia mammillata*.

(B) Pattern of expression of *Eph1* in *Phallusia mammillata* by WMISH.

Expression in *Ciona* at the corresponding stages can be found in the Aniseed database:

*Nodal*: https://www.aniseed.cnrs.fr/aniseed/gene/show_expression?unique_id=Cirobu.g00010576

*Eph1*: https://www.aniseed.cnrs.fr/aniseed/gene/show_expression?unique_id=Cirobu.g00000642

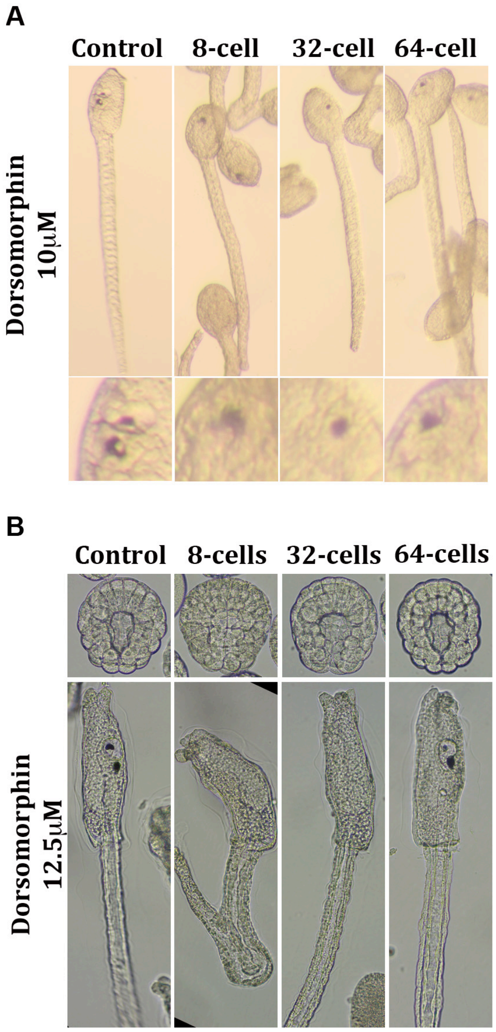


| Treatment with 12.5 μM Dorsomorphin at: | control | 8-cells | 32-cells | 64-cells |
|---|---|---|---|---|
| invaginating embryos | 50/50 | 23/50 | 29/51 | 28/50 |
| larvae with 2 pigmented sensory organs | 50/50 | 0/50 | 4/51 | 35/50 |
| larvae with 1 pigmented sensory organs | 0/50 | 9/50 | 9/51 | 14/50 |
| larvae with 0 pigmented sensory organs | 0/50 | 41/50 | 38/51 | 1/50 |

**Figure S3. Dorsomorphin treatment prevents the formation of the otolith in *Phallusia mammillata***

(A) BMP signalling pharmacological inhibition (Dorsomorphin at 10μM) from 8-cell, 32-cell and 64-cell stages prevents the formation of the otolith pigmented sensory organ in *Phallusia mammillata*. (B) BMP signalling pharmacological inhibition with Dorsomorphin at 12.5μM can abrogate otolith and ocellus formation in *Phallusia mammillata*. It produces some degree of invagination defects but with a lower penetrance than the defects observed in pigmented sensory organ formation.

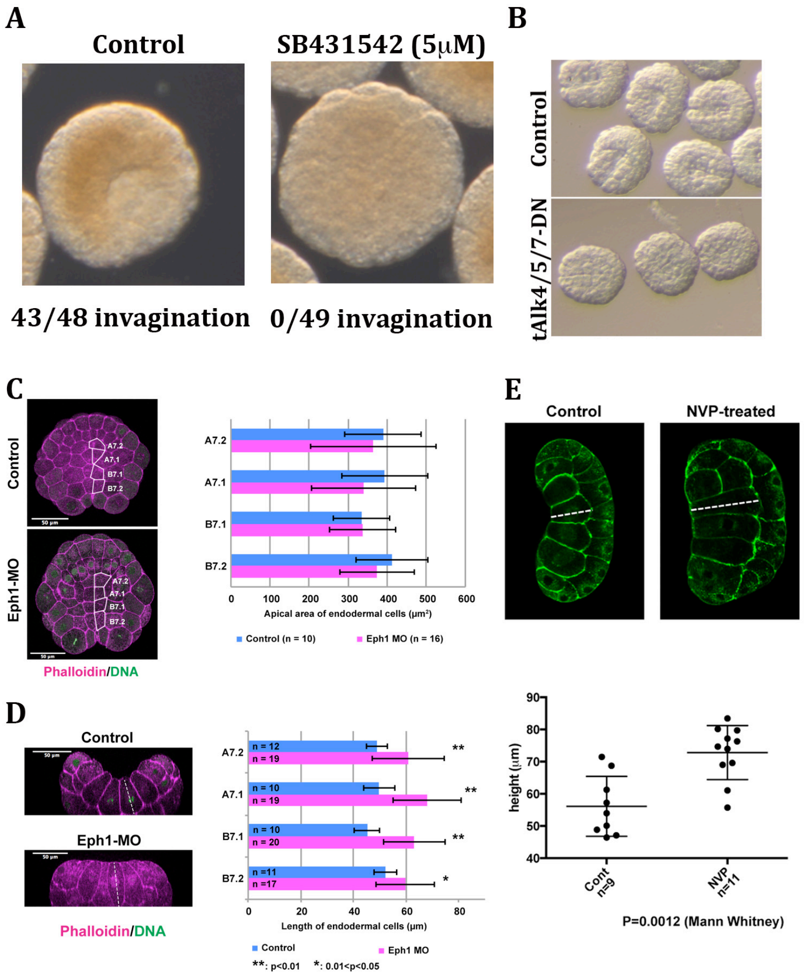


**Figure S4. Inhibition of Nodal and Eph signalling prevents endoderm invagination in *Phallusia mammillata* and *Ciona intestinalis***

(A) Effects on endoderm invagination of treating *Ciona* embryos from 16-cell stage with 5μM SB431542. Results are representative of 2 independent experiments. 43 out of 48 embryos invaginated under control conditions and 0 out of 49 embryos invaginated upon SB431542 treatment (5μM from 16-cell stage). (B) Early gastrula *Phallusia mammillata* embryos microinjected with either FastGreen dye alone (Control) or with mRNA encoding a dominant negative Nodal receptor (tAlk4/5/7). In the controls, 19/20 microinjected embryos invaginated and 15/20 formed a normal larva. 1/17 embryos microinjected with tAlk4/5/7 mRNA invaginated and none formed a larva. (C) Apical area of endodermal *Ciona* cells at the 76-cell stage in control and Eph1-MO-injected embryos. t-test analysis with $p>0.05$ for all blastomeres (D) Height of *Ciona* endodermal cells at the late 112-cell stage (L112-cell) in control and Eph1-MO-injected embryos. The embryos analysed were fixed and stained for actin (Phalloidin) and DNA (DAPI). Statistically significant differences were assessed by t-test analysis (E) Height of *Ciona* endodermal cells at the late 112-cell stage in control and in NVPBHG712-treated (8μM, from 8-cell stage) embryos.

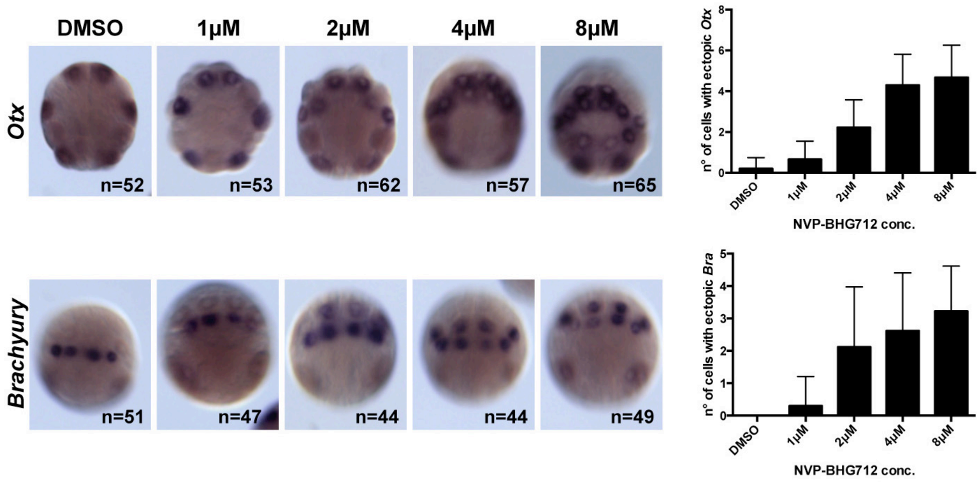


**Figure S5. NVPBHG712 treatment mimics the inhibition of Ephna.d/Eph3 signals**

Treatment of *Ciona* embryos from the 16-cell stage with NVPBHG712 leads to ectopic expression of *Otx* and *Brachyury,* mimicking the effects of Ephna.d/Eph3 inhibition obtained following the microinjection of an *Ephna.d* Morpholino or of a dominant-negative form of the *Eph3* receptor (Picco *et al*, 2007; Ohta and Satou, 2013). The severity of the phenotype is concentration-dependent. NVPBHG712 thus efficiently blocks Eph3 signalling in *Ciona*.

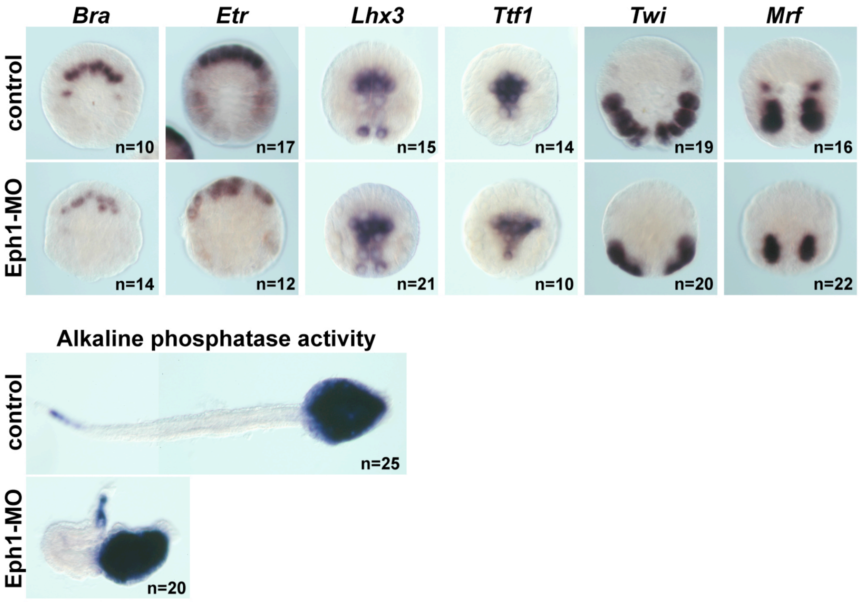


**Figure S6. The major lineage specifications are not affected in Eph1-MO embryos**

Effects of treating *Ciona intestinalis* embryos with Eph1-MO on the major lineage specifications: *Bra* (notochord), *Etr* (neural), *Lhx3* (endoderm), *Ttf1 (endoderm)*, *Twist-like-1* (*Twi*, mesenchyme) and *Mrf* (muscle). (B) Effects of Eph1-MO microinjection on alkaline phosphatase activity (late endoderm marker).

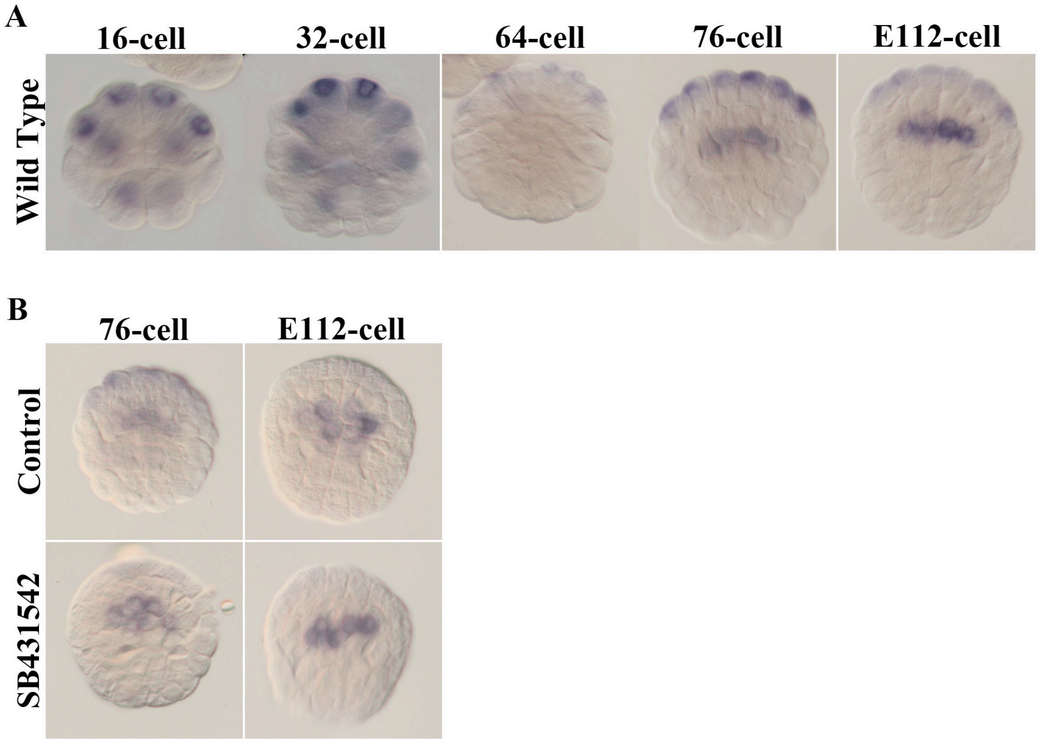


**Figure S7. Nodal signalling inhibition does not affect *Efna.c* expression**

(A) Expression pattern of *Efna.c* by WMISH at 16-cell, 32-cell, 64-cell, 76-cell and early 112-cell (E112 cells) stage in *Phallusia mammillata* embryos. (B) Expression of *Efna.c* in *Phallusia mammillata* embryos under control (DMSO treated) and SB431542 treated (10μM from 16-cell stage) conditions at 76-cell and E112-cell stage. Data representative of 3 independent experiments.

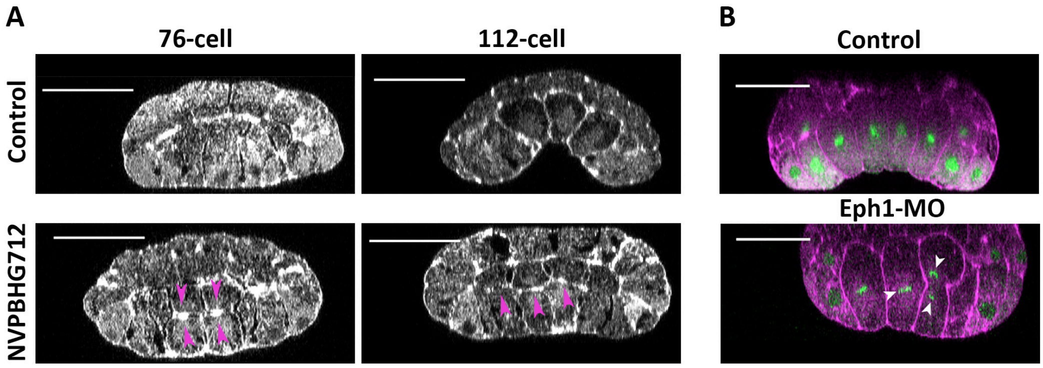


**Figure S8. Eph kinase inhibition and Eph1-MO micro-injection induce premature endoderm cell division** (A) Frontal optical sections through fixed 76-cell stage and late 112-cell stage *Phallusia mammillata* embryos treated with 8 µM NVPBHG712 from the 8-cell stage and stained for actin with Phalloidin. Note the premature division of the endoderm progenitors (arrows). In *Phallusia* 10/64 (15.6%) analysed embryos treated with 8µM NVPBHG712 from the 8-cell stage exhibited premature endoderm division, while none of the control embryos (n=81) showed this phenotype. Scale bar 50 µm (B) *Ciona intestinalis* embryos under control conditions or treated with Eph1-MO. Note the premature division of the endoderm progenitors (arrows). The embryos are stained for actin with Phalloidin and for DNA with DAPI. Scale bar - 50 µm.
